# Bumble bee microbiota shows temporal succession and increase of lactic acid bacteria when exposed to outdoor environments

**DOI:** 10.1101/2023.11.21.568059

**Authors:** Arne Weinhold, Elisabeth Grüner, Alexander Keller

## Abstract

**Question:** The large earth bumble bee (*Bombus terrestris*) maintains a social core gut-microbiota, similar as known from the honey bee, which plays an important role for host health and resistance. Experiments under laboratory conditions with commercial hives are limited to these vertically transmitted microbes and neglect variability by environmental influences and external acquisition of microbes. Various environmental and landscape-level factors may have an impact on the gut-microbiota of pollinating insects, with consequences for pollinator health and fitness in agroecosystems. Still, it is not fully clear whether access to a higher vs lower flower diversity will have a significant influence on the bumble bee microbiota. Here, we tested in a semi-field experiment how strongly the bumble bee microbiota changes over time when exposed to different flower diversities within outdoor flight cages. We used commercial hives to distinguish between vertically and horizontally transmitted bacteria, respectively from the nest environment or the exposed outside environment.

**Result:** The sequential sampling of foraging workers over a period of 35 days indicated a temporal progression of the bumble bee microbiota when exposed to outside conditions. The microbiota became not only more diverse, but changed in composition and variability over time. We observed a major increase in relative abundance of the families *Lactobacillaceae*, *Bifidobacteriaceae* and *Weeksellaceae*. In contrast, major core taxa like *Snodgrassella* and *Gilliamella* declined in their relative abundance over time. The genus *Lactobacillus* showed a high diversity and strain specific turnover, so that only specific ASVs showed an increase over time, while others had a more erratic occurrence pattern. Exposure to different flower diversities had no significant influence on the bumble bee microbiota.

**Conclusion:** The bumble bee microbiota showed a dynamic temporal progression with distinct compositional changes and diversification over time. The exposure of bumble bees to environmental conditions, or environmental microbes, increases dissimilarity and changes the gut-community composition compared to laboratory rearing conditions. This shows the importance of environmental influences on the temporal dynamic and progression of the bumble bee microbiota.

**Scope statement:** Bumble bees (*Bombus terrestris*) are, next to the honey bee, commercially important pollinators and widely used to enhance crop pollination service within greenhouse environments. They host a similar, but characteristic, set of core-microbiota which are of known importance for bumble bee health. Despite this, bumble bees harbor their own specific set of symbionts, which do not occur within the honey bee and seem to be more easily influenced by colonization of environmental microbes. While experiments under controlled lab-based rearing conditions often lack the influence of environmental or landscape-level drivers, field-based observation can often not resolve the influence of a single factor. One major unresolved question is which environmental factor influences the microbiota of social pollinators by environmental microbes. Especially whether monocultures (low flower diversity) are *per se* rather detrimental to microbiota composition compared to more balanced and diverse pollen provisions (high flower diversity). Within this article, we investigated the influence of different flower diversities as potential drivers of the bumble bee gut-microbiota under semi-field conditions. We used outdoor cages which contained a flower diversity gradient to specifically test how a low and high diversity of flower resources could influence the bumble bee microbiota over time.

## 1 Introduction

Bumble bees play an important role for ecosystem service worldwide, due to their role as pollinators for a large variety of plants (Klein et al., 2007; Garibaldi et al., 2013). They are of high commercial value, as they can be used for the pollination of various agricultural-grown plants within field environments (Goulson, 2003; Nayak et al., 2020) and are bred for commercial use in glasshouse environments (Velthuis and Van Doorn, 2006). On some crops, e.g. tomatoes, they are even more effective in pollination than honey bees, due to characteristics like buzz pollination (Vallejo-Marín, 2022), and given the current threats of diseases and parasites as Varroa mites to honey bees, alternative native species are in need to maintain crop and wild plant seed sets (Kevan et al., 1990; Garibaldi et al., 2013; Parreño et al., 2022). To preserve the vital services that bumble bees provide to ecosystems and agriculture, it is essential to prioritize their health and conservation. Especially in agricultural landscapes, increased land use intensity and monocultures cumulate several stressors like pesticides and lowered nutritional quality with negative effects on bumble bee health and colony fitness (Straub et al., 2023). Likewise to other insect groups, bumble bee diversity and abundance has been declining for decades with lower reproduction success in agricultural landscapes compared to urban environments (Williams and Osborne, 2009; Samuelson et al., 2018). Major issues are the reduction in floral resources and diversity of food plants as well as the lack of appropriate nesting sites (Goulson et al., 2008). Additional stressors are the excessive use of pesticides and the introduction of novel pathogens due to international trading (Colla et al., 2006; Stanley and Raine, 2016).

Microbes play an essential role for bee health and resistance, as they help not only with digestion and nutrient uptake (Zheng et al., 2017; Bonilla-Rosso and Engel, 2018), but provide protection against stressors like pathogens, parasites and toxins (Engel et al., 2012; Cariveau et al., 2014; Daisley et al., 2020; Motta et al., 2022). For the large earth bumble bee (*B. terrestris*) as well as the common eastern bumble bee (*B. impatiens*), the microbiota is an important driver for the resistance against infections with the parasite *Crithidia bombi* (Koch and Schmid-Hempel, 2011b, 2012; Mockler et al., 2018). Similar to the honey bee, bumble bees are well known for their simple, but distinct, gut microbiota comprised of a low diversity of characteristic groups belonging to the genera *Snodgrassella* (*Neisseriaceae*), *Gilliamella* (*Orbaceae*), *Lactobacillus* (*Lactobacillaceae*) and *Bifidobacterium* (*Bifidobacteriaceae*) (Koch and Schmid-Hempel, 2011a; Martinson et al., 2011; Powell et al., 2016; Kwong et al., 2017; Hammer et al., 2021a). These groups are considered as corbiculate bee core-bacteria as they are conserved among *Bombus* and *Apis* species (Kwong and Moran, 2016; Raymann and Moran, 2018). Besides these, bumble bees contain *Bombus*-specific groups, which are lacking in honey bees i.e. *Schmidhempelia* (*Orbaceae*) and *Bombiscardovia* (*Bifidobacteriaceae*) (Killer et al., 2010; Martinson et al., 2014).

*Gilliamella* and *Snodgrassella* are known for their complementary metabolic abilities in carbohydrate metabolism (Kwong et al., 2014; Zheng et al., 2019), but showed also a role in parasite protection. A loss of *Snodgrassella* and *Gilliamella* could result in colonies with higher parasite infection rates as well as higher abundance of *Lactobacillus* (Barribeau et al., 2022). While for *Bombus impatiens* a higher abundance of *Apibacter*, *Lactobacillus* and *Gilliamella* spp. was associated with lower pathogen load (Mockler et al., 2018). All those are examples of the crucial roles that a socially transmitted microbiota plays for bee health. Even when reared indoors, bumble bees are able to maintain large parts of their core-microbiota (Meeus et al., 2015). These are maintained through different modes of social transfer and are usually conserved over different life-stages (Billiet et al., 2017b; Su et al., 2021; Zhang and Zheng, 2022). *Snodgrassella* and *Gilliamella* for example are mainly vertically transmitted to the offspring via the queen and are the first microbes to colonize the adult gut (Sauers and Sadd, 2019). Hence, they are not only well preserved within the hive environments, but show high host-specificity as *Snodgrassella* strains from honey bees (*Apis*) cannot colonize bumble bees (*Bombus*) and vice versa (Kwong et al., 2014; Sauers and Sadd, 2019). Each of these symbionts can be split into an *Apis*-specific group (*S. alvi*, *G. apis* or *G. apicola*) as well as a *Bombus*-specific group (*S. communis*, *G. bombicola* or *G. bombi*) (Ludvigsen et al., 2018; Cornet et al., 2022). Another major component of the bee microbiota are ‘lactic acid bacteria’, which are a polyphyletic grouping of Lactobacillales (Firmicutes), and Bifidobacteriales (Actinobacteria) (Olofsson and Vásquez, 2008). These groups are mainly horizontally acquired and require contact to siblings within the nest, while others can also be transmitted by contact to the nesting material (Billiet et al., 2017b).

Besides these hive-maintained core-set of microbes, bumble bees can acquire several strains from the environment, which are considered non-core members, as they are usually lacking in laboratory rearing (Hammer et al., 2021a). Environmental acquisition can have a dominant influence on the microbiota of *B. terrestris* (Bosmans et al., 2018; Krams et al., 2022). A shift in the bumble bee microbiota composition when moved outdoors suggests that particularly enterobacteria are acquired from outdoor environments. Though not considered core-members, enterobacteria can dominate the gut microbiota of bumble bees with up to 90 % relative abundance (Parmentier et al., 2016). During environmental acquisition, flowers could serve as dispersal hubs for beneficial as well as detrimental microbes (Figueroa et al., 2019; Adler et al., 2021; Keller et al., 2021). Thus foraging behavior and available floral sources can have a relevant influence on the microbiota of pollinators (Koch et al., 2012; Newbold et al., 2015; Miller et al., 2019; Martin et al., 2022). Flower species richness and density have been shown to influence bee abundance and are considered as an important aspect for bee health (Doublet et al., 2022). Change of nectar source or pollen availability in agroecosystems could have an influence on the bumble bee microbiota with potentially negative consequences for bumble bee health and resistance. Hence, it is important to better understand how environmental factors and landscape level drivers influence the bumble bee microbiota and which microbial taxa are acquired from the environment. It remained a larger question how much the microbiota is determined by the hosts genetic background, or whether this depends on random exposure to environmental microbes (McFrederick et al., 2012; Näpflin and Schmid-Hempel, 2018).

In this study we examined, how the microbiota of the bumble bee *B. terrestris* changes over time when exposed to outdoor environments. We placed ten bumble bee colonies within a semi-field experiment into separate outdoor flight cages to answer the following questions: (1) How much does the gut-microbiota composition and diversity of adult bumble bees change over time when exposed to outdoor environments? (2) Does the exposure to different flower diversities influence the gut-microbiota of adult bumble bees?

## 2 Material and Methods

### 2.1 Preparation of the field plots

Experiments were conducted in 2022 at the Biocenter of the Faculty of Biology of the Ludwig-Maximilians-University of Munich. We built a total of ten free flight cages using durable and non-impregnated nets as well as pine wood poles that covered a plot area of 2 × 2 meter and 1.75 meter height. Plants that are known to be frequently visited by bumble bees were sown out in eight of the plots in advance to bumble bee hive deposition: *Trifolium repens*, *Trifolium pratense* and *Brassica napus*. To create plots with higher plant diversity, four of the plots included seeds of *Phacelia tanacetifolia*, *Medicago sativa, Borago officinalis* and *Papaver rhoeas*. In each plot 75 g of seeds were used. Two additional plots (9 & 10) were built around already existing native plants which were accessible to native pollinators. If necessary, plots were watered and plant growth observed on a weekly basis. As the first eight plots were built in early April, all plants growing inside were sheltered from visitation of other pollinators. About ten weeks after sowing, the plots were sorted according to the observed flower diversity including also naturally growing plants. Pictures were taken of each plot to index the blooming plants inside, which were ranked from 0 (lowest diversity) to 9 (highest diversity). Despite this planned setup of flower diversity gradient, individual bumble bees managed to escape and foraged on an unknown diversity of flowers outside of the outdoor flight cages.

### 2.2 Bumble bee sampling and sample processing

We obtained large earth bumble bees (*Bombus terrestris*) from a commercial seller (Biobest Group NV, Westerlo, Belgium). Bumble bees were either provided as ‘Mini Hives’ containing about 30 worker bumble bees (plot 1-8) or as ‘Super Mini Hives’ with around 40 workers (plot 9-10). All mini hives were equipped with a care-free nutrition system containing 1.5 liter of sugar solution and pollen supplement to guarantee bumble bee survival during transportation. One hive was placed into each of the plots and covered with cardboard and plastic foil as protection against rain and strong sunshine exposure. Bumble bees were able to leave the mini hive and forage within the flight cages *ad libitum.* The experiments with the bumble bees were conducted under permit: ROB-55.1-8646.NAT_02-8-81-11 according to the nature conservation act of Bavaria (Verordnung zur Ausführung des Bayerischen Naturschutzgesetzes, AVBayNatSchG). Before placement into the plots, one bumble bee from each mini hive was sampled as time point zero (‘t0’). After the placement it took a few days for the bumble bees to adapt to outdoor conditions and actively fly within free flight cages of each plot. As soon as individual bumble bees were seen flying, up to two individuals were sampled per time point and plot. As not all adult bumble bees from every colony were foraging at the same day, we collected some samples over multiple days and binned these for the analysis into seven sampling time points since release in the outdoor flight cages on June 22^nd^ 2023: ‘t0’ (day 0), ‘t1’ (day 13/14), ‘t2’ (day 16/17), ‘t3’ (day 20), ‘t4’ (day 23), ‘t5’ (day 27) ‘t6’ (day 35). On the final sampling day (July 27^th^, 2023), the hive entrances were closed in the early morning, and all animals within the colony immobilized and killed at −20°C. The hives were opened and two adults as well as one larva sampled from inside of each colony. No larvae could be obtained from the hive of plot 2, as there were none inside. Due to vandalism, two of the ten colonies (9 & 10) had to be sampled earlier, so that the final sampling (‘t6’) contains four adults from inside the colony sampled at day 27.

### 2.3 Sample processing, library preparation and sequencing

Frozen bumble bees were dissected using flame sterilized tweezers to obtain the entire gut including crop, foregut and hindgut. For larval samples the entire body was used for DNA isolation. In total, 118 adult guts and 9 larval samples were processed. DNA isolation was performed using the ZymoBIOMICS 96 DNA Kits (Zymo Research) including bead beating at 3200 rpm for 15 min on a grant MPS-1 multiplate shaker (Grant Instruments). Negative extraction controls (NECs) as well as mock-community positive controls (Zymo Research) were included.

We used a dual-indexing approach to amplify the V4 region of the 16S rRNA gene as done by Kozich et al (2013). This protocol includes barcoded primers containing Illumina adapter, index sequence, pad sequence and linker, followed by the gene specific primer 515f 5’-GTGCCAGCMGCCGCGGTAA-3’ and 806r 5’-GGACTACHVGGGTWTCTAAT-3’ (Caporaso et al., 2011). PCR amplification was performed using a Phusion Plus PCR Master Mix (Thermo Scientific) with the following program: 98°C for 30 sec, followed by 30 cycles of 98°C for 10 sec, 55°C for 10 sec, 72°C for 30 sec and a final chain elongation step at 72°C for 5 min. PCR amplification was done in triplicates (3 × 10µl) following the pipetting scheme from (Sickel et al., 2015). PCR products were checked on a E-Gel Power Snap Plus Electrophoresis Device (Thermo Fisher Scientific) using a 96 well E-gel with 1 % Agarose and SYBR Safe. PCR products were normalized using SequalPrep Normalisation Plates (Invitrogen) and pooled into four plate pools. Library quality and fragment size of the plate pools was checked using the High Sensitivity DNA Chip on a 2100 Bioanalyzer (Agilent Technologies). DNA concentration was measured with 1×dsDNA HS Assay Kit on a Qubit 4 Fluorometer (Thermo Fisher Scientific). The four plate pools were pooled equimolarly to a final dilution of 2 nM and paired-end sequenced (2 × 250) on an Illumina MiSeq platform (LMU Biocenter Martinsried) with 5 % PhiX control spiked into the library.

### 2.4 Illumina sequence processing and Microbiota data analysis

To prepare the sequencing data for further analysis, it was processed using VSEARCH v2.14.2 (Rognes et al., 2016) following the metabarcoding processing pipeline available at https://github.com/chiras/metabarcoding_pipeline (Leonhardt et al., 2022). Paired ends of forward and reverse reads were joined, and all reads shorter than 150 bp were removed. Furthermore, quality filtering (EE < 1) as described by Edgar and Flyvbjerg (2015) and *de-novo* chimera filtering following UCHIME3 (Edgar, 2016b) was performed. VSEARCH was also used to define amplicon sequence variants (ASVs) (Edgar, 2016b). By using VSEARCH against the RDP reference database, reads were directly mapped with global alignments with an identity cut-off threshold of 97 %. To classify still remaining reads without taxonomic allocation at this point, SINTAX was used with the same reference database (Edgar, 2016a).

The raw dataset contained 3,887,305 reads and was clustered into 756 ASVs. Non-microbial reads of host organelles like chloroplasts were removed from the dataset. Based on prevalence abundance plots low abundant and low prevalent ASVs were filtered using a quality threshold of 100 reads minimum total abundance and a minimum prevalence of 2 samples within the entire dataset. This step removed in sum only 0.16 % of reads from the *Bombus* samples, but eliminated all extreme low abundant and spurious phyla from the dataset (i.e. Acidobacteria, Armatimonadetes, candidate division WPS−1, Gemmatimonadetes, Planctomycetes, Tenericutes and Verrucomicrobia). The final dataset contained quality ASVs from the phyla Proteobacteria, Firmicutes, Actinobacteria and Bacteroidetes.

Further all ASVs of the mock community used as positive control were filtered from the dataset to account for possible spillover into the samples. Low throughput sample cutoff was set to a minimum of 800 reads per sample (similar as observed for NEC samples). This step removed three larvae and one adult sample with low sequencing throughput from the dataset, retaining bumble bee samples had a median sample sum of 26987 reads (117 adults and 6 larvae). ASVs were binned on genus level and low abundant genera with less than 500 reads total abundance (RA <0.015 %) were removed, filtering 0.06 % of total reads from the dataset. The final dataset contained 116 ASVs of 26 genera. Most of the analysis was performed with the dataset containing only the adult samples.

For the most abundant ASVs obtained the taxonomic assignments were further manually checked against the NCBI Nucleotide Collection and RefSeq Genome Database using nucleotide BLAST (blastn). The closest matching taxa were used together with ASV sequences to construct a phylogenetic tree using the Neighbor-Joining method in MEGA11 to cross-check for a correct phylogenetic placement (Supplemental figure S 1). In this regard, ASV43 was renamed from ‘*Orbus*’ to ‘*Schmidhempelia*’ and ASV11 was renamed from ‘*Bifidobacterium*’ to ‘*Bombiscardovia*’. For ASV6 the taxonomic placement was unclear due to the lack of culturable type strains and closest match to ‘unculturable Firmicutes’ from European bumble bees (Koch and Schmid-Hempel, 2011a). It was renamed from ‘Firmicutes’ to ‘*Xylocopilactobacillus* cf.’ as it seems closely related to recently isolated novel Lactobacillaceae strains from carpenter bees (Kawasaki et al., 2023). While some of the ‘*Snodgrassella*’ and ‘*Gilliamella*’ ASVs were renamed to ‘*Snodgrassella*-like’ and ‘*Gilliamella*-like’ as they indicate a more distant placement with more than 5 % sequence variants to these strains. Percentage identities to *Snodgrassella communis* of 92.94 % (ASV1626), 94.49 % (ASV912) and 94.88 % (ASV863). Percentage identities to *Gilliamella bombi* of 92.13 % (ASV1546), 92.52 % (ASV1536) and 94.9 % (ASV175).

### 2.5 Statistical analysis

R (version 4.3.1) was used for statistical analysis including the ‘phyloseq’ package (McMurdie and Holmes, 2013). The core microbiome was defined with a minimum prevalence of 5 % and minimum relative abundance of 0.1 %. We used linear mixed effect models (lmm) with ‘cage’ as random factor as implemented in the ‘nlme’ package 3.1 (Pinheiro et al., 2023) to investigate the influence of flower diversity or sampling time point on the Shannon diversity. Permutational multivariate analysis of variance using the Bray-Curtis distance matrices (PERMANOVA) was performed as implemented in the adonis2 function with 9999 permutations and sample dissimilarity over time by using the ‘betadisper’ function from the ‘vegan’ package. The influence of sampling time point on the increase and decrease of specific bacterial families and genera was tested by a generalized linear model (glm) using a quasipoisson regression. The obtained p-values from the glm analyses were corrected for multiple testing using the BH method.

## 3 Results

### 3.1 Adult bumble bees show a simple microbiota composition dominated by major core-taxa

We performed a semi-field experiment using outdoor flight cages to investigate how the provision of different flower diversities might change the gut-microbiota of the large earth bumble bee (*B. terrestris*) over time. Adult bees were consecutively sampled within seven sampling time points over a period of 35 days and their gut microbiota analyzed by 16S metabarcoding.

The overall community composition of adult bumble bees showed a relative low diversity and was dominated largely by the families *Neisseriaceae, Orbaceae* and *Lactobacillaceae* (Figure 1A). These families form the major core-microbiota and were found with high prevalence in nearly all individuals. Together with *Bifidobacteriaceae* and *Weeksellaceae* they are responsible for a relative abundance (RA) of 99.3 % of the entire community. Across all samples, the dominating genera were *Snodgrassella* (RA 41.4 %), *Gilliamella* (RA 33.1 %) and *Lactobacillus* (RA 14.7 %). The majority of reads for *Snodgrassella* and *Gilliamella* could be accounted each to a single ASV (Figure 1B), which matched to strains like *S. communis* (ASV1 RA 40.8 %) as well as *G. bombi* (ASV2 RA 32.5 %), both previously isolated from bumble bees (Praet et al., 2017; Cornet et al., 2022) (Supplemental figure S 1). Other *Gilliamella*-like and *Snodgrasella*-like ASVs showed a more distant placement to these type strains, but occurred in rather low abundance. The third most abundant family was *Lactobacillaceae*, which showed overall a high strain diversity with multiple ASVs within the genus *Lactobacillus* (Figure 1B). When applying the phylotype nomenclature used in the past for the honey bee (Ellegaard et al., 2015), these *Lactobacillus* spp. would be accounted to the ‘Firm-5’ clade closely related to *Lactobacillus bombicola*, *L. panisapium* and *L. apis* (Supplemental figure S 1). With 2 % relative abundance *Xylocopilactobacillus* cf. (ASV6) was the second most abundant genus after *Lactobacillus* and represents probably a novel phylotype of bumble bee-related *Lactobacillaceae* (Supplemental figure S 1). Other characteristic *Bombus-*related symbionts were *Bombiscardovia* (RA 1.7 %) (Killer et al., 2010) and *Schmidhempelia* (RA 0.2 %) (Martinson et al., 2014) (Figure 1B). *Apilactobacillus* and *Bombilactobacillus* (‘Firm-4’) showed each with less than 0.07 % only a very low relative abundance.

**Figure 1.**
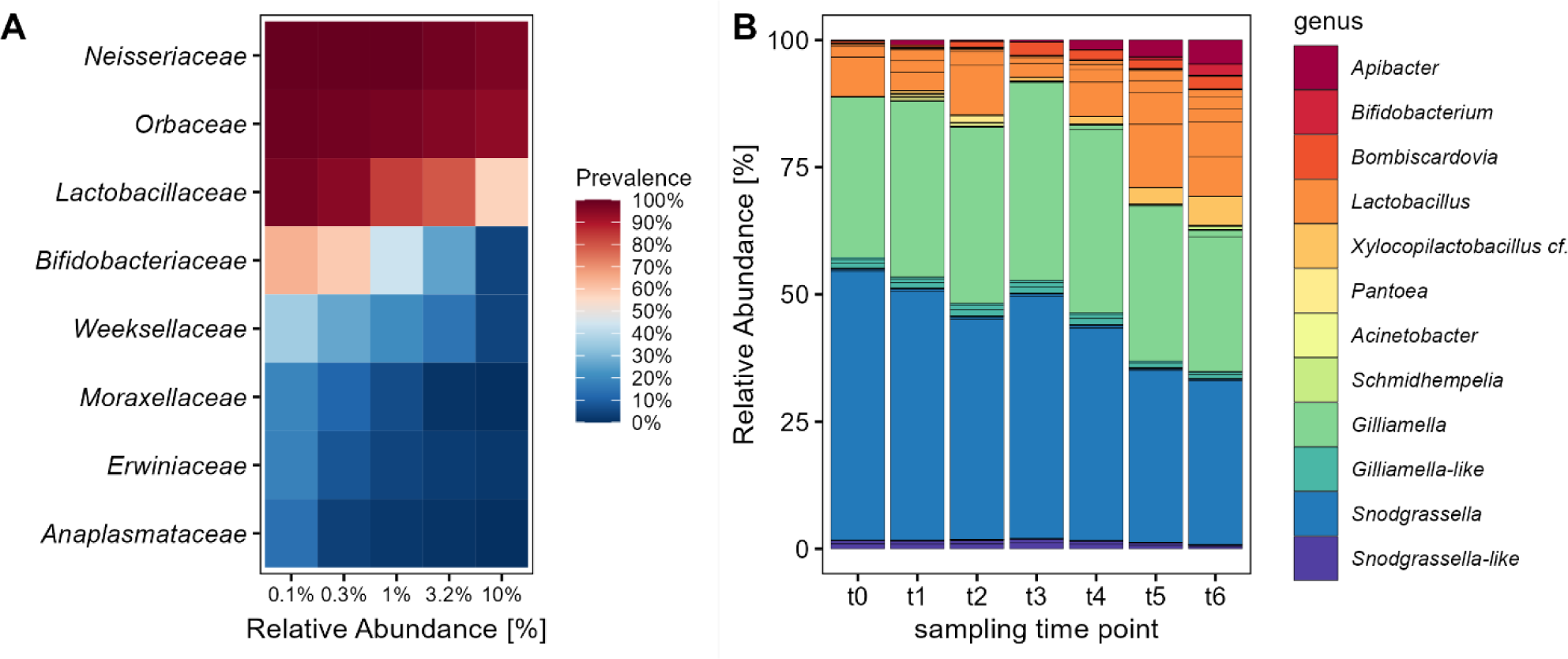
The composition of the large earth bumble bee (*B. terrestris*) gut-microbiota changes over time with a decrease of major core-taxa. (A) Core analysis of the most abundant bacterial families within the gut-microbiota of adult bumble bees across all sampling time points. The families *Neisseriaceae* until *Weeksellaceae* make up to 99.3 % relative abundance. (B) Relative distribution of the bacterial community on ASV level, colored by genus level. Foraging worker of *B. terrestris* were sampled in six sampling time points since release into outdoor flight cages for a period of 35 days. Only bacterial genera with relative abundance of >0.2 % are shown.

### 3.2 Bumble bee microbiota increase in diversity and dissimilarity over time

Despite the simplicity of the bumble bee microbiota the genera *Apibacter*, *Bifidobacterium*, *Bombiscardovia*, *Lactobacillus* and *Xylocopilactobacillus* cf. indicate an increasing relative abundance over the course of the seven sampling time points (Figure 1B). We tested with Linear Mixed-Effects Models with cage as random factor, if there is a temporal change in alpha diversity of the microbial communities and found a significant influence of sampling time point on the Shannon index. Since the release into outdoor flight cages there was a linear increase in alpha diversity on ASV level (lmm: *t* = 5.17, *p* < 0.0001) as well as on genus level (lmm: *t* = 3.73, *p* = 0.0003). This increase in sample diversity was even more pronounced on ASV level (*R*^2^ = 0.19) than on genus level (*R*^2^ = 0.11) (Figure 2). In addition, we tested whether the provision of different flower diversities within the different flight cages would influence the bumble bee microbiota. There was no linear correlation between flower diversity and diversity of the bumble bee microbiota on ASV level (lmm: *t* = −1.149, *p* = 0.284) nor on genus level (lmm: *t* = −0.167, *p* = 0.871) (Supplemental figure S 2A,B). Reasons for the lack of an effect within this setup is discussed later.

**Figure 2.**
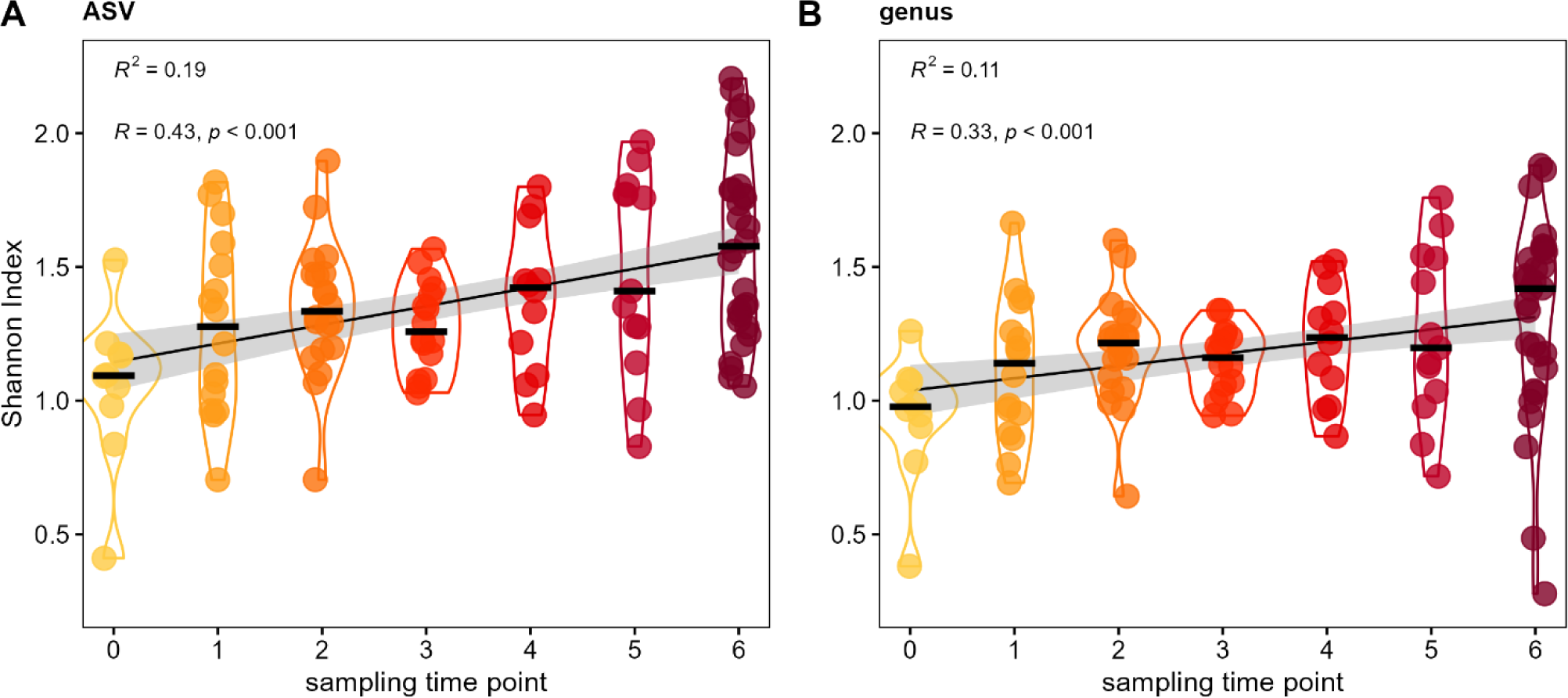
Diversity of the bumble bee gut-microbiota increases by sampling time point. Temporal increase in Shannon diversity on ASV level (A), as well as genus level (B). Foraging bumble bees (*B. terrestris*) were sampled in different sampling time points (‘t0’ to ‘t6’) since release into outdoor flight cages.

Besides this temporal progression of alpha diversity increase, we investigated whether dissimilarity among individual samples would also change over time, i.e. whether individuals from different colonies become more different to each other. Beta diversity was shown by Bray-Curtis distance using non-metric multidimensional scaling (NMDS) colored by sampling time point (Figure 3A). To better illustrate the temporal changes, each time point is shown and highlighted in an individual plot from the same NMDS (Figure 3B-H). Sampling time point had a significant influence on the Bray-Curtis distance (PERMANOVA *F*_1,116_ = 13.99, *p* < 0.001). Beta diversity expanded particularly in the last two sampling time points (‘t5’ and ‘t6’), which showed the highest sample dissimilarity within the dataset (Figure 3G,H). By applying a mixed effects model, community dissimilarity changes significantly over time independent from colony identity (lmm: *t* = 5.07, *p* < 0.0001) (Figure 3I). The largest differences in beta distance were evident between time point ‘t3’ and ‘t6’ (Wilcoxon test with BH correction *p* < 0.0001). These results show a temporal increase in sample variation so that the microbiota of bumble bees become more diverse over time when exposed to outdoor environments.

**Figure 3.**
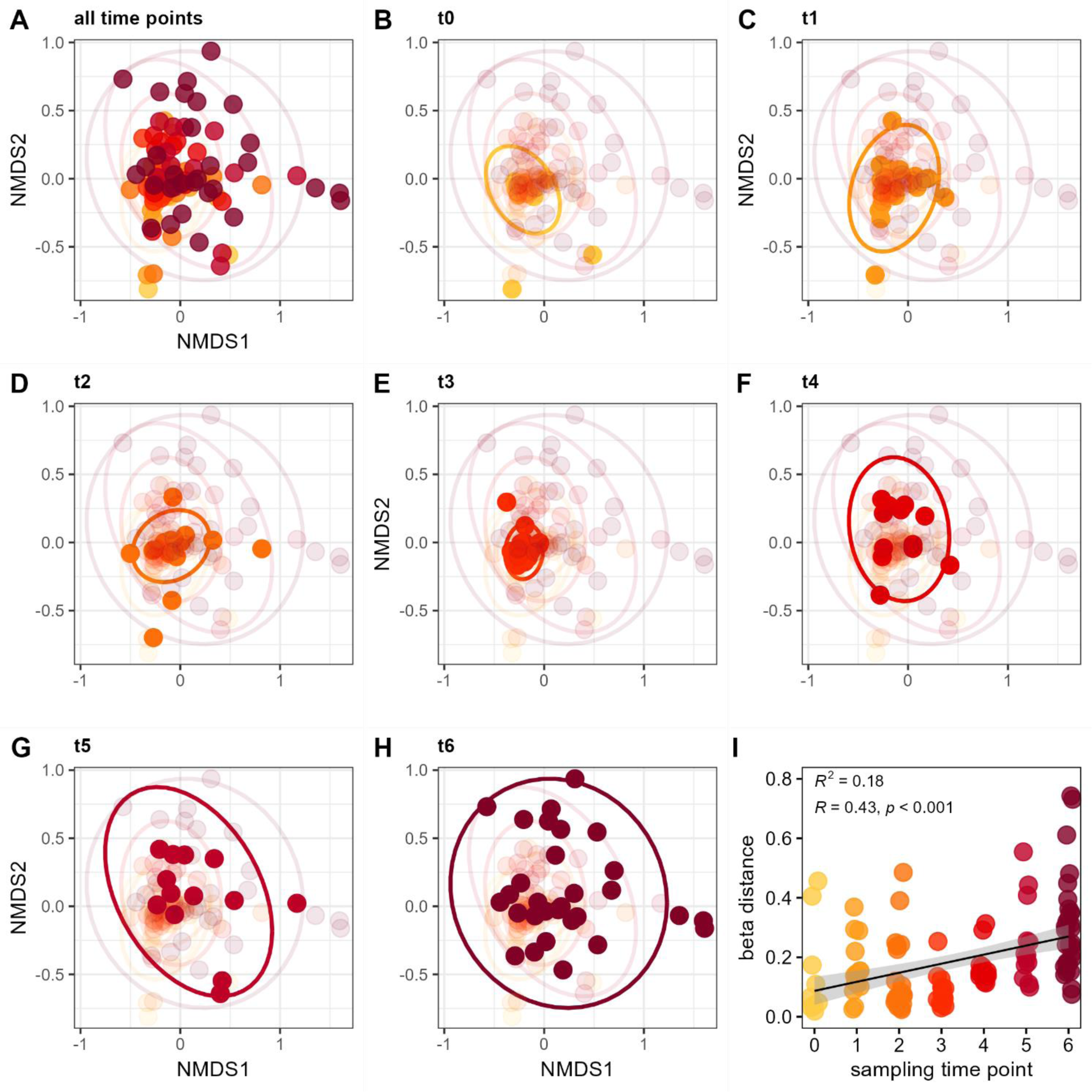
Diversification of the bumble bee gut-microbiota over time. NMDS plots show Bray-Curtis distance for all sampling time points (A), as well as for individual sampling time points ‘t0’ (B), ‘t1’ (C), ‘t2’ (D), ‘t3’ (E), ‘t4’ (F), ‘t5’ (G) and ‘t6’ (H). Increase of beta distance by sampling time points (I). The different time points (‘t0’ to ‘t6’) are indicated by color (yellow to red). Late sampling time points show a higher dissimilarity of the bumble bee microbiota since release into outdoor flight cages.

When applying a similar analysis using food plant provision, we found no influence of flower diversity on microbial community composition (PERMANOVA *F*_9,116_ = 1.31, *p* = 0.15) (Supplemental ^f^igure S ^2^C). Likewise, flower diversity had no significant effect on beta distance (lmm: t = −1.01, *p* = 0.343) (Supplemental figure S 2D).

### 3.3 Temporal turnover of individual bacterial families

To further evaluate which bacterial groups were responsible for the increase in diversity and dissimilarity over time, we looked at the temporal changes in relative abundance of individual bacterial families. This showed that the families of *Bifidobacteriaceae*, *Weeksellaceae* and particularly *Lactobacillaceae* indicate an increase in relative abundance, while *Neisseriaceae* and *Orbaceae* tend to decrease (Figure 4). We used generalized linear models with quasi-poisson distribution and corrected p-values for multiple testing by the BH method. Here we found a positive influence of sampling time point on the relative abundance of *Bifidobacteriaceae* (glm: *t* = 4.81, *p* < 0.0001), *Weeksellaceae* (glm: *t* = 2.76, *p* = 0.01) and *Lactobacillaceae* (glm: *t* = 4.85, *p* < 0.0001). The latter showed such a drastic increase that some bumble bee samples from the final sampling time point (‘t6’) were even dominated by *Lactobacillaceae* (Figure 4). On the other hand, there was a reciprocal trend for other families to decrease in relative abundance. The core-families *Neisseriaceae* (glm: *t* = −5.63, *p* < 0.0001) and *Orbaceae* (glm: *t* = −2.23, *p* = 0.034) showed a significant decrease in their relative abundance over the course of the sampling period (Figure 4). Others, like the family of *Erwiniaceae* showed no temporal trend over time (glm: *t* = −1.75, *p* = 0.082), but occurred only occasionally in a few samples with low relative abundance in the entire dataset (RA <0.4 %). This shows that the temporal diversification of the bumble bee microbiota was mainly due to an increase in relative abundance of the families *Bifidobacteriaceae*, *Weeksellaceae* and *Lactobacillaceae*, while the abundance of major core-members within the *Neisseriaceae* and *Orbaceae* decreased.

**Figure 4.**
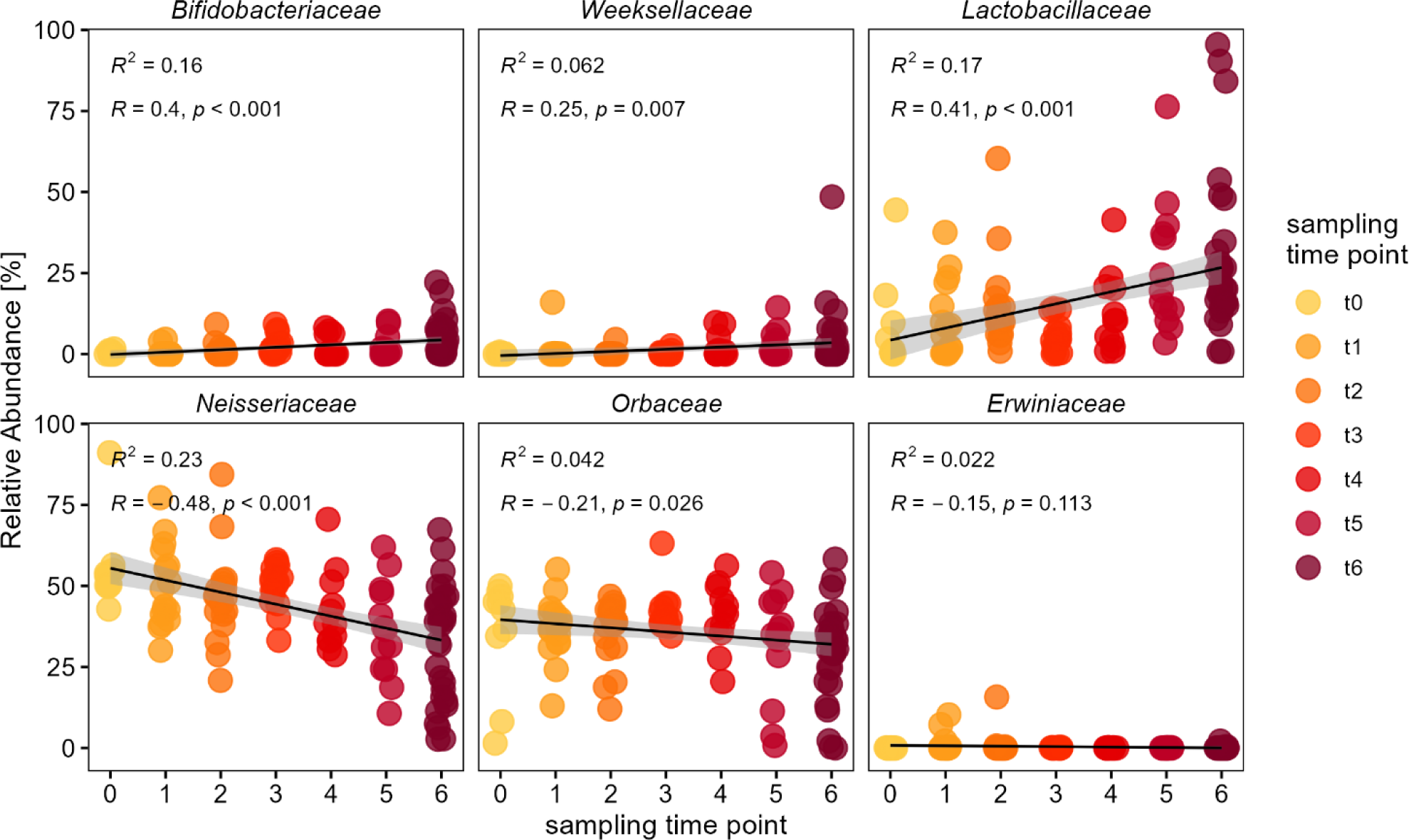
Temporal change in relative abundance of individual bacterial families within the bumble bee gut-microbiota. Relative abundances of individual bacterial families since release into outdoor flight cages. The families *Bifidobacteriaceae*, *Weeksellaceae* and *Lactobacillaceae* show an increase in relative abundance, while major core-taxa i.e. *Neisseriaceae* and *Orbaceae* show a decrease over time. The different sampling time points (‘t0’ to ‘t6’) are indicated by color (yellow to red). Only major families with a cumulative relative abundance of 99.7 % are shown.

### 3.4 Temporal progression on genus level

For a more detailed analysis we also investigated temporal changes of the most abundant bacterial genera (Figure 5). *Apibacter* was the only genus among the *Weeksellaceae* and showed the same pattern on genus level (glm: *t* = 2.76, *p* = 0.01). Among the *Bifidobacteriaceae*, both genera of *Bifidobacterium* (glm: *t* = 2.96, *p* < 0.01) as well as *Bombiscardovia* (glm:, *t* = 2.81, *p* < 0.01) showed a significant increase in relative abundance over time. In the family *Lactobacillaceae* the genera of *Lactobacillus* (glm: *t* = 3.61, *p* = 0.0012) as well as *Xylocopilactobacillus* cf. (glm: *t* = 4.29, *p* < 0.001) showed an increase in relative abundance over time (Figure 5). The family *Neisseriaceae* showed the strongest trend for a temporal decrease mainly due to a significant decrease of the genus *Snodgrassella* (glm: *t* = −5.40, *p* < 0.0001), as well as for the low abundant *Snodgrassella*-like ASVs (glm: *t* = −4.07, *p* < 0.001). Though overall more variable in abundance, the family of *Orbaceae* showed still a significant decrease of the genus *Gilliamella* (glm: *t* = −2.15, *p* = 0.04) as well as for the *Gilliamella*-like ASVs (glm: *t* = −3.58, *p* = 0.001), but not for *Schmidhempelia* (glm: *t* = 0.54, *p* = 0.59).

**Figure 5.**
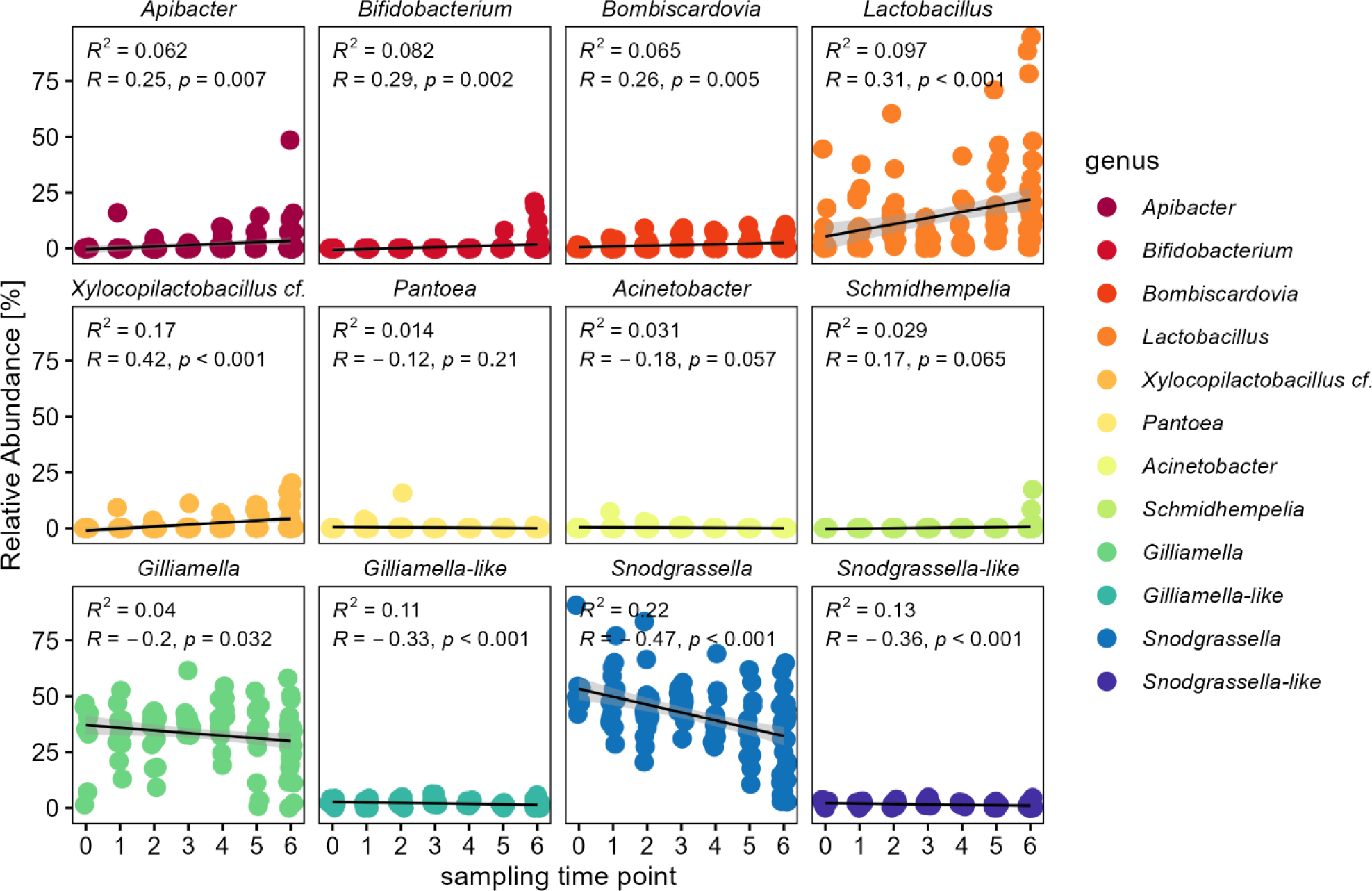
Temporal change in relative abundance of individual bacterial genera within the bumble bee gut-microbiota. Relative abundances of individual genera show an increase of *Apibacter* (*Weeksellaceae*), *Bifidobacterium* and *Bombiscardovia* (*Bifidobacteriaceae*), *Lactobacillus* and *Xylocopilactobacillus* cf. (*Lactobacillaceae*). Major core-taxa show a decrease in relative abundance: *Gilliamella* (*Orbaceae*) and *Snodgrassella* (*Neisseriaceae*). Only genera with relative abundance of >0.1 % are shown.

### 3.5 Comparison of adults and larvae from the final sampling time point

At the final sampling time point (‘t6’) bumble bees were not only sampled outside of the colonies by a net, but as well from inside the colony. For this analysis we included the few larval samples (n=6) which have been obtained from the opened hives. We found only marginal differences in community composition among the sampling groups from the final time point (PERMANOVA_t6_ : *F*_2,35_ = 1.93, *p* = 0.042). The adults sampled outside of the colony seem to contain larger abundances of *Apibacter* (RA 8.5 %) compared to the adults sampled from inside the colony (RA 2.0 %), while those from inside the colony showed higher abundance of *Bifidobacterium* (RA 3.8 % vs 0.04 %) (Supplemental figure S 3). Notably, *Schmidhempelia* was only detected in four individuals sampled from inside the colony (4 of 20), but not in any of the foraging adults sampled outside of the colonies (0 of 97). The larval samples differed mainly from the adults as they contained larger relative abundance of *Pediococcus* (RA 16.7 %), which was nearly absent in the adults (RA 0.25 %).

### 3.6 Turnover of individual ASVs among *Lactobacillaceae*

Within the bumble bee microbiota, the family of *Lactobacillaceae* stood out as it contained a much higher ASV diversity compared to other bacterial families. We were interested whether these ASVs show a turnover in their abundance over the sampling time points and if only particular ASVs increase in abundance while other might even decrease. As we compared all major ASVs to the closest matching type strains (Supplemental figure S 1) we were able to obtain a near species level resolution among *Lactobacillus* spp. This allows us to have a deeper look into the dynamics within the family of *Lactobacillaceae* from time point ‘t0’ to ‘t6’ (Supplemental figure S 4). The observed increase in the genus *Lactobacillus* was mainly due to an increase in ASV7 related to *Lactobacillus apis* (glm: *t* = 4.56, *p* < 0.001) as well as ASV5 and ASV26 related to *L. panisapium* (ASV5, glm: *t* = 3.24, *p* < 0.005; ASV26, glm: *t* = 2.31, *p* = 0.051). While those ASVs related to *L. bombicola* showed a more variable abundance over time with no clear trend for an increase (ASV3, glm: *t* = −1.06, *p* = 0.435; ASV4, glm: *t* = 1.12, *p* = 0.435). Hence, the increase within the genus *Lactobacillus* is highly strain specific and only some ASVs within this group show a similar temporal trend, while others have a more erratic occurrence pattern (Supplemental figure S 4). Even on ASV level *Xylocopilactobacillus* cf. (ASV6, glm: *t* = 4.30, *p* < 0.001) shows a significant increase over time and reaches up to 5.9 % RA in the final sampling time point. Other low abundant groups like *Bombilactobacillus* (ASV64, glm: *t* = 0.18, *p* = 0.854) or *Fructobacillus* (ASV55, glm: *t* = −0.32, p = 0.846) indicated no significant change.

## 4 Discussion

### 4.1 Environmental influence and plasticity of the bumble bee microbiota

We investigated how the exposure to outdoor environments changes the microbiota of the bumble bee *B. terrestris*. We found a temporal succession of the bumble bee microbiota with an increase in diversity and sample dissimilarity over time. The bumble bee microbiota in our dataset showed overall a low diversity and was mainly dominated by the genera *Snodgrassella, Gilliamella* and *Lactobacillus* (Figure 1). These are typical core-groups which could be found in most of our individuals and are known to be highly conserved among social corbiculate bees (Kwong and Moran, 2016; Kwong et al., 2017; Zhang and Zheng, 2022). We could demonstrate that the bumble bee microbiota shows a temporal succession with a reduction of prominent core-members *Snodgrassella* and *Gilliamella*, which were replaced mainly by an increasing relative abundance of *Lactobacillus* (Figure 5). Such a shifted microbiota composition has been previously associated with higher parasite infection rates (Barribeau et al., 2022), but it remains unclear whether community shifts are a result of the infections or would render colonies more susceptible. But following the progression of bee microbiota assembly on a temporal gradient has only been investigated in a few studies, i.e. with *A. cerana* (Dong et al., 2021) or *B. impatiens* (Hammer et al., 2023a). Temporal shifts in community composition can be explained by an accumulation of a higher diversity of environmentally acquired strains, so that other core-members appear to diminish in relative abundance.

Even for the bumble bee *B. terrestris* with a socially maintained core-microbiota, environmental influences can have a large impact on the microbial community composition (Newbold et al., 2015; Parmentier et al., 2016). In general, mainly *Enterobacteriaceae*, *Apibacter* (*Weeksellaceae*) and *Fructobacillus* (*Lactobacillaceae*) are considered as environmentally acquired strains, as these groups usually lack in laboratory environments (Newbold et al., 2015; Hammer et al., 2021a). Environmental influences can be shown by location or habitat dependence, as colonies of *B. terrestris* near forest environments were dominated by *Fructobacillus* compared to colonies in agricultural or horticultural landscapes (Krams et al., 2022). An investigation of 28 Chinese bumble bee species revealed two distinct enterotypes either dominated by core-members of the microbiota (*Snodgrassella* and *Gilliamella*) or by externally acquired microbes mainly belonging to *Enterobacteriaceae* (Li et al., 2015). When moving colonies of *B. terrestris* outdoors, the microbiota can shift towards an increase in *Enterobacteriaceae* (Parmentier et al., 2016). Such a shift in wild bumble bee microbiota is often considered as a ‘disrupted’ microbiome and associated with higher pathogen load (Villabona et al., 2023). Overall, the influence of environmental microbes differs a lot between different studies, and it remains unclear what causes such community shifts. Within our dataset, Enterobacteriales showed only a very low abundance and did not contribute to the progression in compositional turnover over time. We observed only an occasional occurrence of *Pantoea* (*Erwiniaceae*) in some of the early time points (RA <0.4 %). Similar, *Acinetobacter* (*Moraxellaceae*) showed only an occasional occurrence with very low abundance (RA 0.2 %), but is a common isolate of honey bees as well as floral nectar (Kim et al., 2014; Alvarez-Perez et al., 2021). Although it is putatively environmentally acquired, *Apibacter* can be considered as typical member of the bumble bee gut-microbiota (Praet et al., 2016; Hammer et al., 2021a; Steele and Moran, 2021). We observed an increase in relative abundance of *Apibacter* over time, similar as shown for the Asian honey bee *A. cerana* (Dong et al., 2021). We also found lower abundance of *Apibacter* in adults sampled from inside the colony compared to foraging adults, which is evidence that this group is mainly environmentally acquired.

### 4.2 Increase and high strain diversity of *Lactobacillaceae*

Similar as for honey bees (Ellegaard et al., 2015), we observed a high diversity of *Lactobacillus* strains in *B. terrestris*. Lactobacilli are a highly diverse group and multiple strains have been isolated from honeybees (Olofsson et al., 2014) as well as other wild bees and flowers (McFrederick et al., 2018). Several of these strains which have been previously classified as ‘*Lactobacillus* spp.’ showed diverging properties and have been later split into different genera (Zheng et al., 2020). These are: *Apilactobacillus* (previously known as the *L. kunkeei* group), *Bombilactobacillus* (previously known as *L. bombi* ‘Firm-4’ group) and *Lactobacillus* (previously known as ‘Firm-5’ group). Here, we would add *Xylocopilactobacillus* cf. as a novel bumble bee associated phylotype. This is probably a novel group of bumble bee-related *Lactobacillaceae* with yet unclear taxonomic placement (distinct from *Lactobacillus*, *Bombilactobacillus* and *Apilactobacillus*) (Supplemental figure S 1). Similar strains have been already cloned from *B. terrestris* in earlier studies (Mohr and Tebbe, 2006) (GenBank: AJ880198), but could not be further classified and were described until now only as ‘uncultured Firmicutes’ from bumble bees (GenBank: HM215045) (Koch and Schmid-Hempel, 2011a). This group has been occasionally reported as ‘Firm-3’ cluster (McFrederick et al., 2013; Leonhardt and Kaltenpoth, 2014) and seems to be characteristic for European bumble bee populations, as it has not been described for *B. impatiens* (Mockler et al., 2018)(Hammer et al., 2023a). This provides opportunities to characterize a new phylotype of *Bombus*-associated Lactobacilli. So far, related culturable strains have only recently been isolated from carpenter bees and characterized as strictly anaerobic with auxotrophy for NAD biosynthesis (Kawasaki et al., 2023). They were proposed as a new genus of *Xylocopilactobacillus* gen. nov (Kawasaki et al., 2023). Although carpenter bees (*Xylocopa*) are not eusocial (but rather facultatively, incipiently or sub-social), their microbiota shows surprising parallels to that of *Bombus* species, with similar conserved core-taxa including *Schmidhempelia, Bombilactobacillus* and *Bombiscardovia* (Gu et al., 2023; Handy et al., 2023). Here it can be speculated that the long life expectancy of the females in *Xylocopa* species which share the nests with the offspring adult generation (Velthuis and Gerling, 1983), allows for a similar microbial transfer as otherwise only known from eusocial corbiculate bees.

For bumble bees, the relationship with lactic acid bacteria seems to be highly strain specific (McFrederick et al., 2013) and adults usually require the direct contact to nestmates for an acquisition and propagation of this group within the hive (Billiet et al., 2017b). *B. terrestris* cannot be colonized by generic *Lactobacillus* strains as a probiotic treatment, while *Bombus*-specific strains showed stable colonization (Billiet et al., 2017a). This shows that bee-related *Lactobacillus* strains cannot be replaced by other generic strains. The proliferation and diversification of lactic acid bacteria within bumble bee guts point at an important functional role of this group for host fitness. Lactic acid bacteria are known for their importance to honey bee health (Vásquez et al., 2012; Killer et al., 2014; Iorizzo et al., 2022) and resemble an important part of the bumble bee microbiota. For some ground nesting bees they can be even the dominating taxon within their gut-microbiota (Hammer et al., 2023b).

In our dataset, the genus *Lactobacillus* showed a high strain diversity on ASV level, which further proliferated across the sampling time points. The temporal increase in this genus could be mainly observed for the strain *Lactobacillus apis* (ASV7), originally isolated from honey bees (Killer et al., 2014), as well as *L. panisapium* (ASV5, ASV26) isolated from bee bread (Wang et al., 2018). This could be indication that these groups have been acquired via direct or indirect contact with honey bees during bumble bee foraging. Other *Lactobacillus* ASVs were related to *L. bombicola* (ASV3, ASV4), which had been previously described from bumble bees (Praet et al., 2015). These showed a more erratic occurrence within individual bumble bee samples with no clear temporal trend towards an in- or decrease in abundance. Whether this means that this strain might be hive-maintained and is not environmentally acquired is not fully clear.

As an alternative explanation, environmental temperatures could influence community composition in bumble bees when exposed to outdoor conditions. An increase in rearing temperatures had a positive effect on the proliferation of *Lactobacillaceae* within the gut microbiota of *B. impatiens* (Palmer-Young et al., 2019). Hence, even putative *Bombus*-specific strains like *Xylocopilactobacillus* cf. could proliferate in their relative abundance due to increasing temperatures without the need for an acquisition from environmental sources. However, the core taxa *Snodgrassella* and *Gilliamella* show likewise a better growth rate at elevated temperatures (Hammer et al., 2021b), but were decreasing in relative abundance within the course of our sampling period.

Behavioral experiments with *B. impatiens* showed that bumble bees seem to avoid flowers inoculated with *Apilactobacillus micheneri,* pointing at a deterring effect of some lactic acid bacteria from environmental sources (Russell and Ashman, 2019). This strain was previously isolated as *Lactobacillus micheneri* from the gut of sweat bees *Halictus ligatus* and has been associated with flowers and other megachilid bees (McFrederick et al., 2017, 2018). In contrast, the inoculation of nectar with *Fructobacillus* lead to an increased nectar consumption by *B. impatiens* (Russell and McFrederick, 2022). For solitary bees, which do not exchange microbes via social contact, environmental acquisition from flowers is often the only source to obtain a more diverse microbiota (Voulgari-Kokota et al., 2019a, 2019b; Cohen et al., 2020).

### 4.3 Temporal shifts of the bumble bee microbiota

The microbiota of bees can show dynamic plasticity over time, when followed over different life stages and seasons (Dong et al., 2021; Li et al., 2021; Su et al., 2021). For *B. terrestris*, developmental changes have been investigated for different larval stages, which differed clearly in their microbiota compared to the adults (Guo et al., 2023). Larvae of *B. terrestris* have been described to be mainly colonized by *Lactobacillus* (Su et al., 2021), while we found all major core groups from the adults within the larvae. The major difference was the colonization by an unspecific *Pediococcus* (*Lactobacillaceae*). But the overall lower sequencing depth in our larval samples is also indicative for a much lower microbial biomass in the larvae compared to the adults. As a result, three of the nine larval samples needed to be removed due to low sequencing depth. Upon hatching, adult bumble bees, much like honeybees, emerge bacteria-free and acquire their microbiota from their food, hive environment or nestmates (Koch and Schmid-Hempel, 2011b; Hammer et al., 2021b). This process happens within the first 4 days of the adult life so that the overall microbial load remains relatively stable with progressing adult age for *B. impatiens* (Hammer et al., 2023a). When reared indoors, the microbiota of *B. terrestris* and *B. impatiens* shows no larger change in alpha diversity over time (Parmentier et al., 2016; Hammer et al., 2023a). This was clearly different in our setup, as the placement into outdoor environments resulted in diversification of bumble bee microbiota, observable by an increase in in alpha diversity as well as an increase in sample dissimilarity over time. Especially the increase in dissimilarity from time point ‘t4’ to ‘t6’ could indicate that a new generation of worker have emerged into a more diverse hive environment.

Though diversity levels did not change, Hammer *et al*. (2023a) reported a community shift of the bumble bee microbiota with age, resulting in a decrease in *Schmidhempelia* and the establishment of *Gilliamella*, while proportions of *Lactobacillus* remain relatively stable over a period of 60 days. Though *Schmidhempelia* has been described as dominant member of the microbiota of the common eastern bumble bee (*B. impatiens*) (Hammer et al., 2023a), we found it only with low abundances within a few individuals of *B. terrestris*. We observed also larger shifts in community composition, but a decrease in relative abundance of *Gilliamella*, while *Lactobacillaceae* were increasing within a 35 day period. Here, it is important to note that the previous study with *B. impatiens* was conducted in a laboratory setting, whereas our study used *B. terrestris* was performed under environmental conditions in outdoor cages. Seasonal changes and sampling time point are also strong predictor of the honeybee microbiota independent from their geographic location (Almeida et al., 2023).

### 4.4 Why flower diversity had no influence on the bumble bee microbiota

There are several possible explanations why flower diversity of the provided food plants had no significant influence on the bumble bee microbiota within our setup (Supplemental figure S 2). First, only a few of the sowed plants bloomed early enough to provide nectar and pollen in sufficient quantities so that the bumble bees relied primarily on the resources provided by their mini hives. Hence, the provided flower density might have been too low to show an effect. Second, our initial setup excluded other pollinators and does not allow visitation and cross-species transfer of microbes from wild pollinators (but only wind-dispersed microbes). Here, it would be interesting to further elucidate whether increased plant diversity alone, or only in combination with a broader range of pollinating insects might yield a different outcome. At least, floral diversity has an influence on pollinator diversity, so that both factors are difficult to disentangle (Doublet et al., 2022). As the third reason, several bumble bees manage to escape through tiny holes that have been bitten into the nets and could be observed returning from foraging flights outside of the cages. Hence, they were exposed to an unknown diversity of flowering plants outside of the assigned area and could introduce microbes from the surrounding environment. Even though they showed an excellent sense of orientation and returned precisely to their specific hives, this all blurs the influence of the provided flower diversity gradient. As a result, the ten cages with the treatment groups did not differ in their microbial diversity nor dissimilarity and conclusions about flower diversity should be taken with caution.

While social transfer is the most important route for bumble bees to maintain a conserved core-microbiota, floral visitation provides further chances for microbial acquisition and transfer (Miller et al., 2019), but increases also the risk of pathogen exposure from other pollinators (Davis et al., 2021) (Nicholls et al., 2022). Hence the maintenance of a social core that protects bumble bees during their first flights from parasite infections is of great importance. Still, they are able to acquire a more diverse microbiota from their surrounding environment. Bumble bees are even well capable of dispersing microbes among flowers themselves, as demonstrated with *B. impatiens* (Russell et al., 2019). Here, flowers should not only be seen as a source of food provision, but as well as dispersal hubs for environmental microbes, so that vectoring insects move microbes along the plant-pollinator network (McFrederick et al., 2017; Keller et al., 2021; Zemenick et al., 2021; Weinhold, 2022).

## 6 Supplemental Figures

**Supplemental figure S 1.**
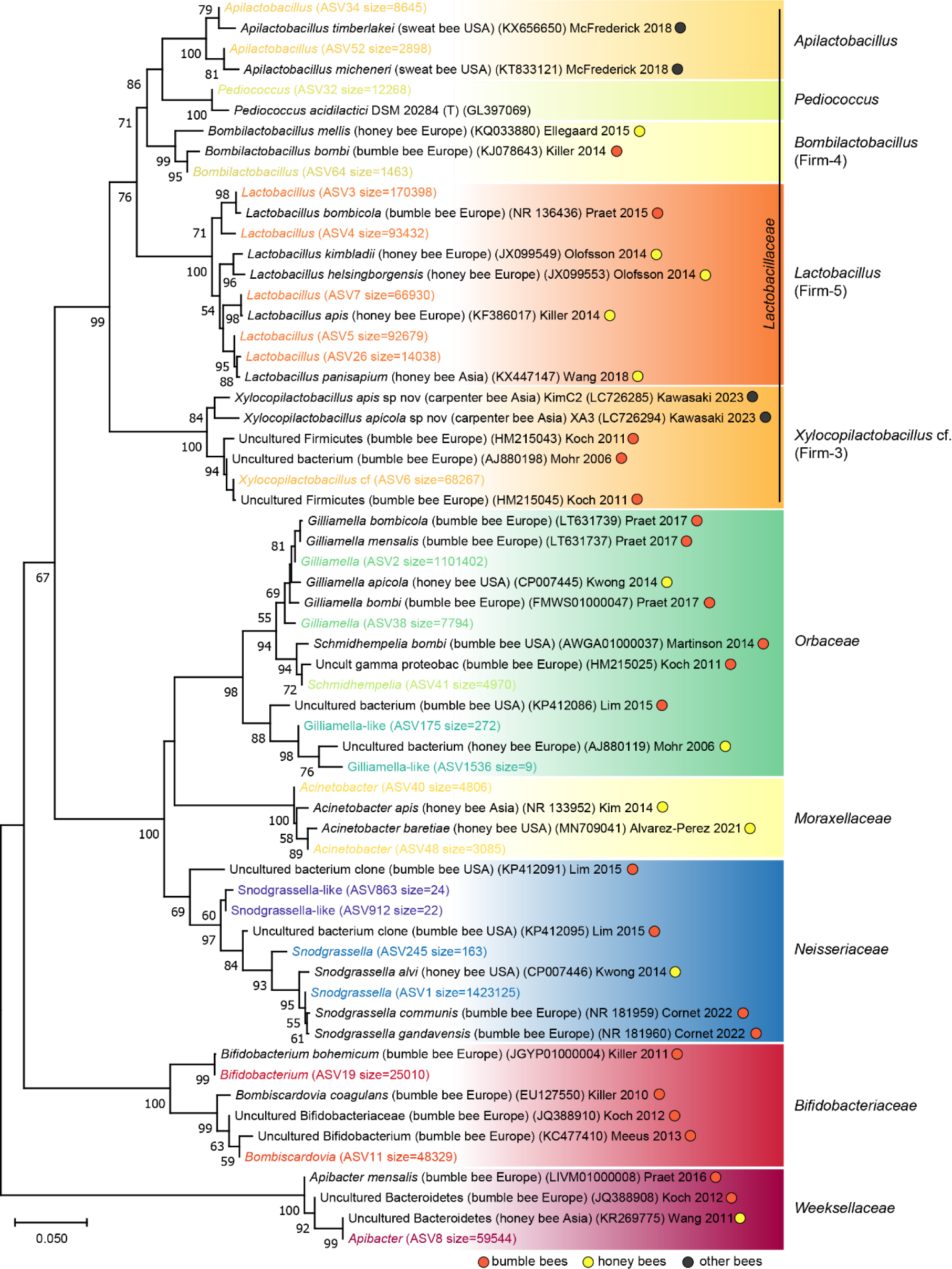
Phylogenetic tree of major ASVs obtained from the bumble bee gut-microbiota (*B. terrestris*). Bumble bee ASVs (in color) were aligned to the closest matching sequences obtained from the NCBI Nucleotide Collection database. Isolation source, geographic origin and references are indicated for each sequence. Neighbor-Joining tree was constructed with MEGA11 and bootstrapping values >50 with 1000 repetitions are shown next to the branches.

**Supplemental figure S 2.**
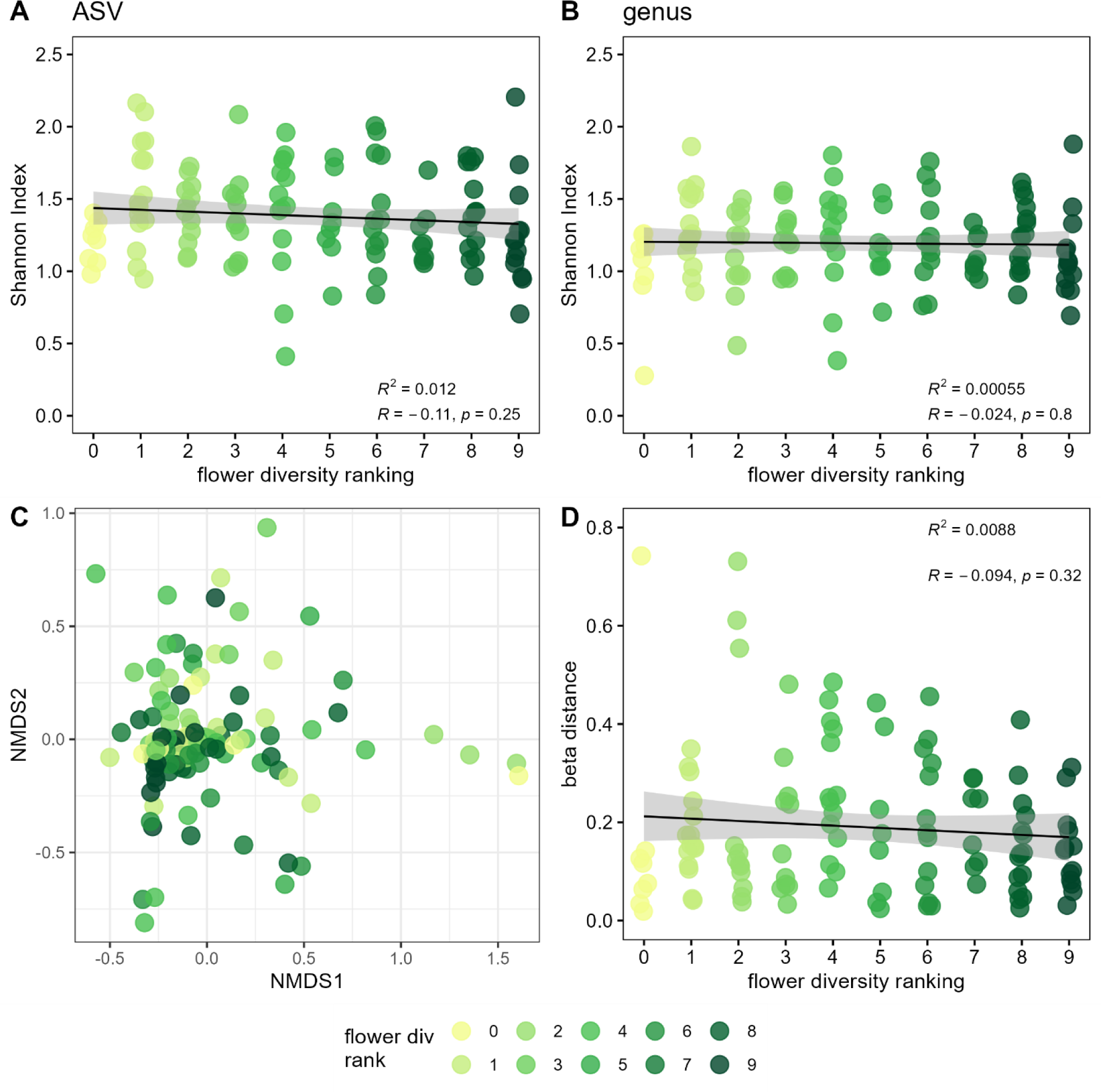
Food plant diversity had no influence on the gut-microbiota diversity of the large earth bumble bee (*B. terrestris*). Shannon diversity on ASV (A) and genus level (B) shown by food plant diversity ranking. Bray-Curtis distance of microbial communities shown as NMDS plot (C) and beta distance (D) colored by food plant diversity. Bumble bee worker were sampled from colonies reared in ten individual outdoor flight cages ranked from low to high food plant diversity (0 to 9).

**Supplemental figure S 3.**
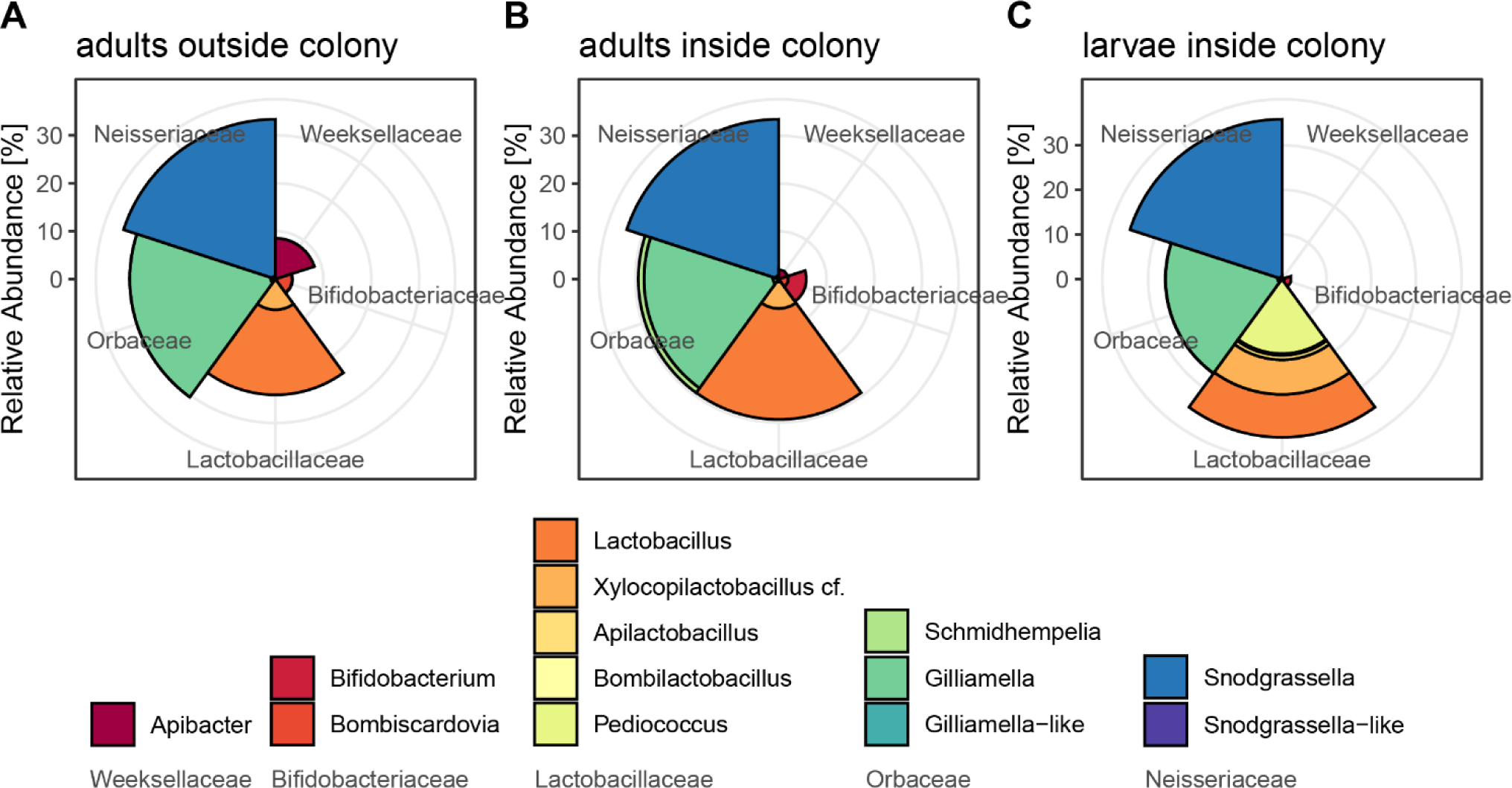
Comparison of the bumble bee gut-microbiota of foraging worker with adults and larvae sampled from inside the colony. All samples (adults outside colony, adults inside colony, larvae inside colony) were taken at the final sampling time point (t6). Foraging adults indicate higher abundance of *Apibacter*, but lower abundance of *Bifidobacterium*. *Schmidhempelia* was only detected in adults sampled from inside the colony. Larval samples show larger abundance of *Pediococcus*.

**Supplemental figure S 4.**
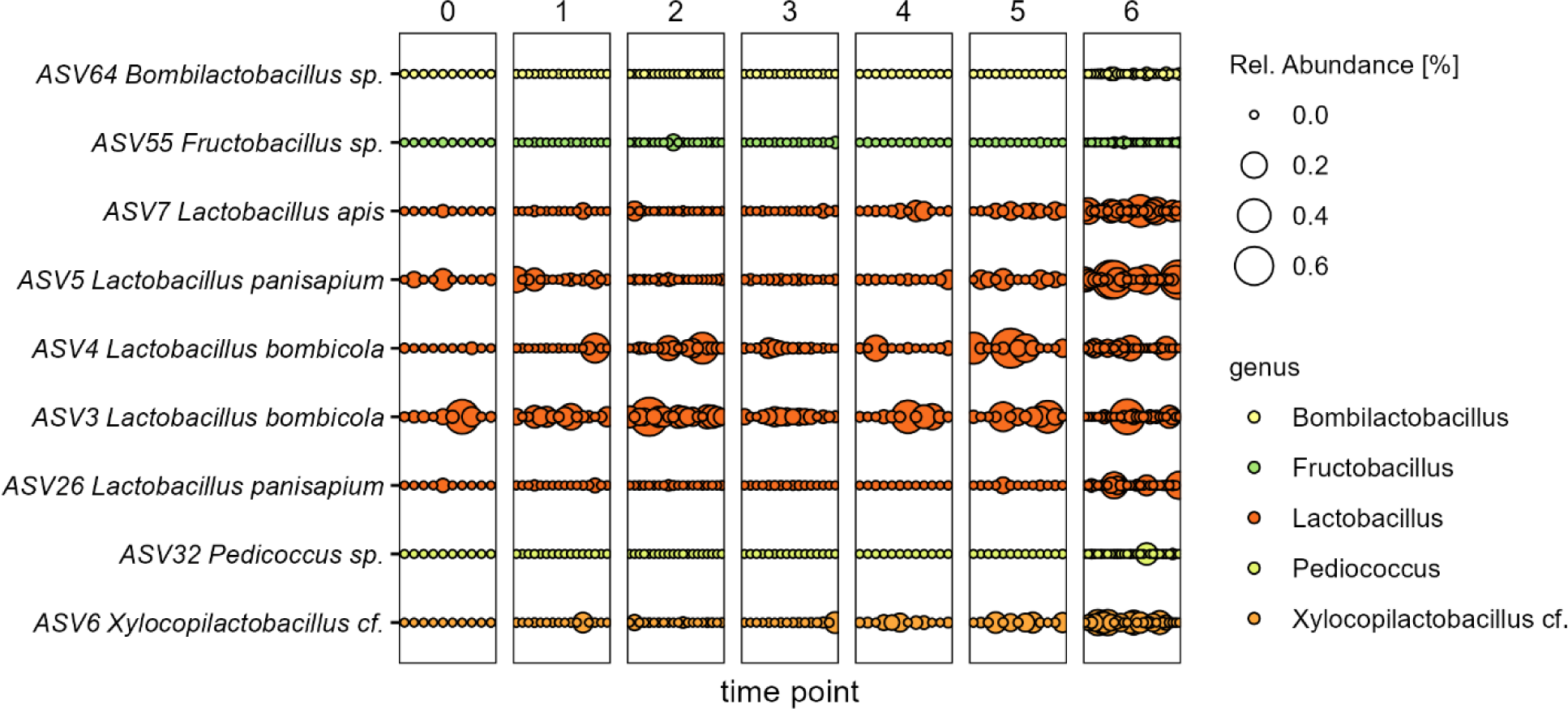
ASV turnover within *Lactobacillaceae* over time. Nine most abundant ASVs within the *Lactobacillaceae* are shown in their abundance dynamic within individual samples from time point ‘t0’ to ‘t6’. Detailed taxonomy of ASVs obtained from Supplemental figure S 1. Increase in relative abundance over time can be primarily observed for *Lactobacillus apis* (ASV7), *L. panisapium* (ASV5, ASV26) and *Xylocopilactobacillus* cf. (ASV6). Other strains like *L. bombicola* (ASV3, ASV4) show a more erratic and variable abundance with no clear temporal trend.

## 7 Conflict of Interest

The authors declare that the research was conducted in the absence of any commercial or financial relationships that could be construed as a potential conflict of interest.

## 8 Author Contributions

AW: Conceptualization, Formal analysis, Data curation, Investigation, Methodology, Visualization, Statistical analysis, Writing – original draft, Writing – review and editing; EG: Conceptualization, Formal analysis, Investigation, Methodology, Visualization, Writing – review and editing; AK: Conceptualization, Funding acquisition, Project administration, Resources, Supervision, Writing – review and editing.

## 9 Funding

AW acknowledges support by the LMUexcellent Postdoc Support Fund.

## 10 Acknowledgments

The authors would like to thank Lars Landgraf for help during insect sampling, Anna Preußner for help during building of the experimental setup and Uschi Schkölziger for technical assistance. Further, the authors want to thank Andreas Brachmann from the Genomics Service Unit of the LMU for help during sequencing and the government of upper Bavaria for providing sampling permits for *B. terrestris*.

## 11 Data Availability Statement

The dataset generated for this study can be found in the NCBI Sequence Read Archive (SRA) under BioProject number PRJNA1042966. Metabarcoding processing pipeline is available at github: https://github.com/chiras/metabarcoding_pipeline.

## Notes

### Competing Interest Statement

The authors have declared no competing interest.

